# CIAlign - A highly customisable command line tool to clean, interpret and visualise multiple sequence alignments

**DOI:** 10.1101/2020.09.14.291484

**Authors:** Charlotte Tumescheit, Andrew E. Firth, Katherine Brown

**Affiliations:** Department of Pathology, University of Cambridge, Cambridge, UK

**Keywords:** Multiple sequence alignment, alignment quality, Python tool, comparative genomics, transcriptomics, phylogenetics

## Abstract

**Background:** Throughout biology, multiple sequence alignments (MSAs) form the basis of much investigation into biological features and relationships. These alignments are at the heart of many bioinformatics analyses. However, sequences in MSAs are often incomplete or very divergent, which leads to poorly aligned regions or large gaps in alignments. This slows down computation and can impact conclusions without being biologically relevant. Therefore, cleaning the alignment by removing these regions can substantially improve analyses. Manual editing of MSAs is very widespread but is time-consuming and difficult to reproduce.

**Results:** We present a comprehensive, user-friendly MSA trimming tool with multiple visualisation options. Our highly customisable command line tool aims to give intervention power to the user by offering various options, and outputs graphical representations of the alignment before and after processing to give the user a clear overview of what has been removed.

The main functionalities of the tool include removing regions of low coverage due to insertions, removing gaps, cropping poorly aligned sequence ends and removing sequences that are too divergent or too short. The thresholds for each function can be specified by the user and parameters can be adjusted to each individual MSA. CIAlign is designed with an emphasis on solving specific and common alignment problems and on providing transparency to the user.

**Conclusion:** CIAlign effectively removes problematic regions and sequences from MSAs and provides novel visualisation options. This tool can be used to refine alignments for further analysis and processing. The tool is aimed at anyone who wishes to automatically clean up parts of an MSA and those requiring a new, accessible way of visualising large MSAs.

## Introduction

Throughout biology, multiple sequence alignments (MSAs) of DNA, RNA or amino acid sequences are often the basis of investigation into biological features and relationships. Applications of MSAs include, but are not limited to transcriptome analysis, in which transcripts may need to be aligned to genes; RNA structure prediction, in which an MSA improves results significantly compared to predictions based on single sequences; and phylogenetics, where trees are usually created based on MSAs. There are many more applications of MSA at a gene, transcript and genome level involved in a huge variety of traditional and new approaches to genetics and genomics, many of which could benefit from the tool presented here.

An MSA typically represents three or more DNA, RNA or amino acid sequences, which represent partial or complete gene, transcript, protein or genome sequences. These sequences are aligned by inserting gaps between residues to bring more similar residues (either based on simple sequence similarity or an evolutionary model) into the same column, allowing insertions, deletions and differences in sequence length to be taken into account [1, 2]. The first widely used automated method for generating MSAs was CLUSTAL [2] and more recent versions of this tool are still in use today, along with tools such as MUSCLE [3], MAFFT [4], T-Coffee [5] and many more. The majority of tools are based upon various heuristics used to optimise progressive sequence alignment using a dynamic programming based algorithm such as the Needleman-Wunsch algorithm [6]. It has been shown previously that removing divergent regions from an MSA improves the resulting phylogenetic tree [7]. Various tools are available to remove or improve poorly aligned columns, including trimAl [8], Gblocks [7] and various refinement methods incorporated into alignment software [3, 4]. Some tree building software can also take into account certain discrepancies in the alignment, for example RaXML [9] can account for missing data in some columns and check for duplicate sequence names and gap-only columns; similarly GUI based toolkits for molecular biology such as MEGA [10] sometimes have options to delete or ignore columns containing gaps. However, several common issues affect the speed, complexity and reliability of specific downstream analyses but are not addressed by existing tools.

Clean and Interpret Alignments (`CIAlign`) is primarily intended to address four issues which are commonly encountered when working with MSAs. Researchers in many fields regularly edit MSAs by hand to address these issues, however as well as being extremely time consuming, ensuring reproducibility with this approach is almost impossible and it cannot be incorporated into an automated analysis pipeline.

The first issue we intend to address is that it is common for an MSA to contain more gaps towards either end than in the body of the alignment. This problem occurs at both the sequencing and alignment stage. For example, the ends of *de novo* assembled transcripts tend to have lower read coverage [11] and therefore have a higher probability of mis-assembly and therefore mis-alignment. MSAs created using these sequences therefore also have regions of lower reliability towards either end. Similarly, both Sanger sequences and sequences generated with Oxford Nanopore’s long read sequencing technology, which are often used directly in MSAs, tend to have lower quality scores at the either the beginning or the end [12, 13, 14]. Automated removal of these regions from MSAs would therefore increase the reliability of downstream analyses. Also, while generating an MSA, terminal gaps complicate analysis, and the weighting of terminal gaps relative to internal gap opening and gap extension penalties can make a large difference to the resulting alignment [15]. This again leads to regions of ambiguity and therefore gaps towards the ends of the alignment.

Secondly, insertions or other stretches of sequence can be present in a minority of sequences in an MSA, leading to large gaps in the remaining sequences. For example, alignments of sections of bacterial genomes often result in long gaps representing genes which are absent in the majority of species. These gaps can be observed, for example, in multiple genome alignments shown in Tettelin et al. 2005 [16] for *Streptococcus agalactiae* and Hu et al. 2011 [17] for *Burkholderia*, amongst others, which show many genes which are present in only a few genomes. While these regions are of interest in themselves and certainly should not be excluded from all further analysis, they are not relevant for every downstream analysis. For example, a consensus sequence for these bacteria would exclude these regions and their presence would increase the time required for phylogenetic analysis without necessarily adding any additional information. Large gaps in some sequences may also result from missing data, rather than true biological differences and, if this is known to be the case, it is often appropriate to remove these regions before performing phylogenetic analysis [18].

Thirdly, one or a few highly divergent sequences can heavily disrupt the alignment and therefore complicate downstream analysis. It is very common for an MSA to include one or a few outlier sequences which do not align well with the majority of the alignment. One example of this is metagenomic analyses identifying novel sequences in large numbers of datasets. It is common to manually remove phylogenetic outliers which are unlikely to truly represent members of a group of interest (see for example [19–21]) but this is not feasible when processing large numbers of alignments.

Finally, very short partially overlapping sequences cannot always be reliably aligned using standard global alignment algorithms. It is very common to remove these sequences, manually or otherwise, prior to further analysis.

There are also several common issues in alignment visualisation. Large alignments can be difficult to visualise and a small and concise but accurate visualisation can be useful when presenting results, so this has been incorporated into the software. With many alignment trimming tools it can be difficult to track exactly which changes the software has made, so a visual output showing these changes could be helpful.

Finally, transparency is often an issue with bioinformatics software, with poor reporting of exactly how a file has been processed [22–24]. CIAlign has been developed to process alignments in a transparent manner, to allow the user to clearly and reproducibly report their methodology.

CIAlign is freely available at github.com/KatyBrown/CIAlign.

## Materials and Methods

CIAlign is a command line tool implemented in Python 3. It can be installed either via pip3 or from GitHub and is independent of the operating system. It has been designed to enable the user to remove specific issues from an MSA, to visualise the MSA (including a markup file showing which regions and sequences have been removed), and to interpret the MSA in several ways. CIAlign works on nucleotide or amino acids alignments and will detect which of these is provided. A log file is generated to show exactly which sequences and positions have been removed from the alignment and why they were removed. Users can then adjust the software parameters according to their needs.

CIAlign takes as its input any pre-computed MSA in FASTA format containing at least three sequences. Most MSAs created with standard alignment software will be of an appropriate scale, for example single or multi-gene alignments and whole genome alignments for many microbial species. Measurements on the runtime were conducted for MSAs created by randomly drawing equally probable nucleotides and adding gap regions such that each MSA has a certain proportion of gaps. When running CIAlign with all core functions (cleaning functions and creating mini alignments for input, output and the markup) and for fixed gap proportions, the runtime scales quadratically with the size of the MSA, i.e. with *n* as the number of sequences and *m* the length of the MSA, the worst case time complexity is *O*((*nm*)2). Further runtime measurements were taken for running CIAlign with the core functions on an MSA of constant size with different numbers of gaps. The runtime decreases linearly with an increasing proportion of gaps. It should be noted that, besides the size of the MSA and its gap content, the runtime is impacted by which combination of functions is applied. For very long MSAs the size of the final image becomes a limiting factor when creating a sequence logo, as the matplotlib library [25] has restrictions on the number of pixels in one object. We have provided detailed instructions about this limit in the “Guidelines for using CIAlign” on the CIAlign GitHub.

The path to the alignment file is the only mandatory parameter. Every function is run only if specified in the parameters and many function-specific parameters allow options to be fine-tuned. Using the parameter option *--all* will turn on all the available functions and run them with the default parameters, unless otherwise specified. Additionally, the user can provide parameters via a configuration file instead of via the command line.

CIAlign has been designed to maximise usability, reproducibility and reliability. The code is written to be as readable as possible and all functions are fully documented. All functions are covered by unit tests. CIAlign is freely available, open source and fully version controlled.

### Cleaning Alignments

CIAlign consists of several functions to clean an MSA by removing commonly encountered alignment issues. All of these functions are optional and can be fine-tuned using user parameters. All parameters have default values. The available functions are presented here in the order they are executed by the program. The order can have a direct impact on the results, the functions removing positions that lead to the greatest disruptions in the MSA should be run first as they potentially make removing more positions unnecessary and therefore keep processing to a minimum. For example, divergent sequences often contain many insertions compared to the consensus, so removing these sequences first reduces the number of insertions which need to be removed. Sequences can be made shorter during processing with CIAlign and therefore too short sequences are removed last.

Fig 1 shows a graphical representation of an example toy alignment before (Fig 1A) and after (Fig 1B-1F) using each function individually. The remove gap only function is run by default after every cleaning step, unless otherwise specified by the user.

**Fig 1.**
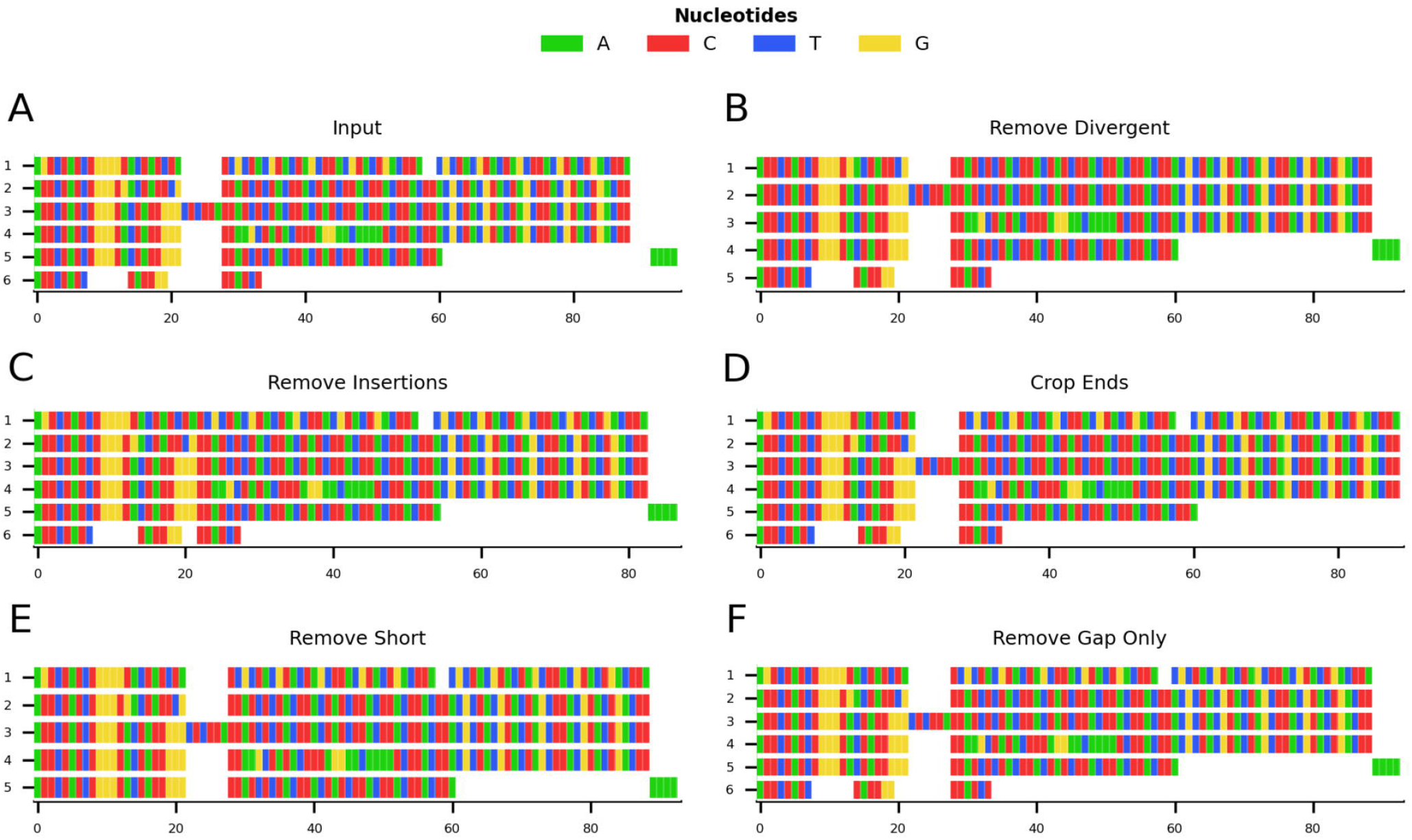
Mini alignments showing the main functionalities of CIAlign based on Example 1. **a** Input alignment before application of CIAlign, generated using the command “ `CIAlign --infile example1.fasta --plot_input`”. **b** Output alignment showing the functionality of the remove divergent function, generated using the command “ `CIAlign --infile example1.fasta --remove_divergent --plot_output`”. **c** Output alignment showing the functionality of the remove insertions function, generated using the command “`CIAlign --infile example1.fasta --remove_insertions --plot_output`”. **d** Output alignment showing the functionality of the crop ends function, generated using the command “`CIAlign --infile example1.fasta --crop_ends --plot_output`”. **e** Output alignment showing the functionality of the remove short sequences function, generated using the command “`CIAlign --infile example1.fasta --remove_short --plot_output`”. **f** Output alignment showing the functionality of the remove gap only function, generated using the command “`CIAlign --infile example1.fasta --plot_output`”. Subplots were generated using the`drawMiniAlignment` function of CIAlign.

#### Remove Divergent

For each column in the alignment, this function finds the most common nucleotide or amino acid and generates a temporary consensus sequence. Each sequence is then compared individually to this consensus sequence. Sequences which match the consensus at a proportion of positions less than a user-defined threshold (default 0.65) are excluded from the alignment (Fig 1B). It is recommended to run the `make_similarity_matrix` function to calculate pairwise similarity before removing divergent sequences, in order to adjust the parameter value for more or less divergent alignments.

#### Remove Insertions

In order to define a region as an insertion, an alignment gap must be present in the majority of sequences and flanked by a minimum number of non-gap positions on either side, which can be defined by the user (default 5). The minimum and maximum size of insertion to be removed can also be defined by the user (default 3 and 200 respectively) (Fig 1C).

#### Crop Ends

Crop ends redefines where each sequence starts and ends, based on the ratio of the numbers of gap and non-gap positions observed up to a given position in the sequence. It then replaces all non-gap positions before and after the redefined start and end, respectively, with gaps. This will be described for redefining the sequence start, however crop ends is also applied to the reverse of the sequence to redefine the sequence end.

The number of gap positions separating every two consecutive non-gap positions is compared to a threshold and if that difference is higher than the threshold, the start of the sequence will be reset to that position. This threshold is defined as a proportion of the total sequence length, excluding gaps, and can be defined by the user (default: 0.05) (Fig 1D, Fig 2).

**Fig 2.**
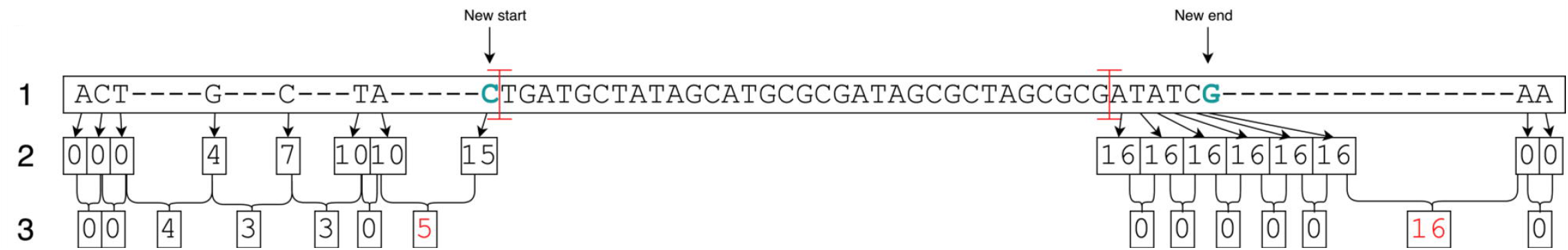
Crop ends diagram. This manually created example illustrates how crop_ends works internally. The length of the sequence shown is 111 including gaps and 80 excluding gaps (1). With a threshold of 10% for the proportion of non-gap positions to consider for change in end positions, 8 positions at the start and at the end, respectively, are being considered (illustrated by red crossbars). For each of these, the number of preceding gaps is calculated (2). Then the change in gap numbers (3) for every two consecutive non-gap positions is compared to the gap number change threshold, which is 5%, i.e. 4 gaps, as a default value. Looking at the change in gap numbers, the last change at each end equal to or bigger than the threshold is coloured in red. This leads to redefining the start and the end of this example sequence to be where the nucleotides are coloured in green.

The user can set a parameter that defines the maximum proportion of the sequence for which to consider the change in gap positions (default: 0.1) and therefore the innermost position at which the start or end of the sequence may be redefined. It is recommended to set this parameter no higher than 0.1, since even if there are a large number of gap positions beyond this point, this is unlikely to be the result of incomplete sequences (Fig 2).

#### Remove short sequences

Remove short sequences removes sequences which have less than a specified number of non-gap positions, which can be set by the user (default: 50) (Fig 1E).

#### Remove gap only columns

Remove gap only removes columns that contain only gaps. These could be introduced by manual editing of the MSA before using CIAlign or by running the functions above (Fig 1F). The main purpose of the function is to clean the gap only columns that are likely to be introduced after running any of the cleaning functions.

### Visualisation

There are several ways of visualising the alignment, which both allow the user to interpret the alignment and clearly show which positions and sequences CIAlign has removed. CIAlign can also be used simply to visualise an alignment, without running any of the cleaning functions. All visualisations can be output as publication ready image files.

#### Mini Alignments

CIAlign provides functionality to generate mini alignments, in which an MSA is visualised using coloured rectangles on a single *x* and *y* axis, with each rectangle representing a single nucleotide or amino acid (e.g. Fig 1, Figs 3-5). Even for large alignments, this function provides a visualisation that can be easily viewed and interpreted. Many properties of the resulting file (dimensions, DPI, file type) are parameterised. In order to minimise the memory and time required to generate the mini alignments, the matplotlib imshow function [25] for displaying images is used. Briefly, each position in each sequence in the alignment forms a single pixel in an image object and a custom dictionary is used to assign colours. The image object is then stretched to fit the axes.

**Fig 3.**
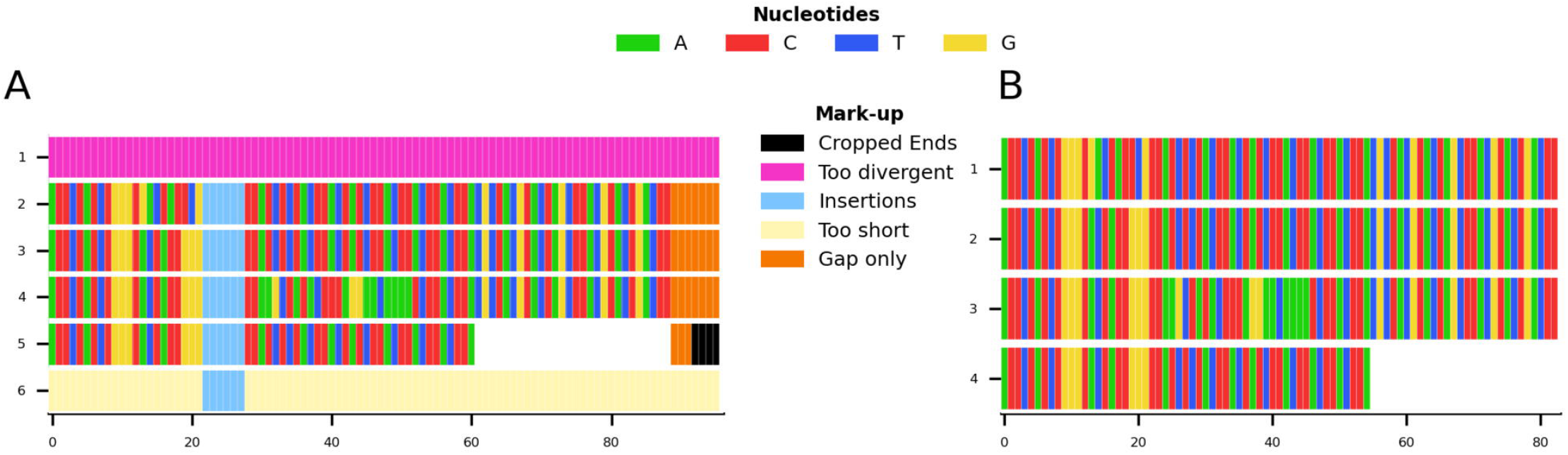
Mini alignments and legends showing further functionalities of CIAlign based on Example 1. **a** Alignment showing the functionality of the plot markup function, generated using the command “`CIAlign --infile example1.fasta --all`”. The areas that have been removed are marked up in different colours, each corresponding to a certain function of CIAlign. **b** Output alignment after application of all functions of CIAlign combined, generated using the command “`CIAlign --infile example1.fasta --all`”. Subplots were generated using the `drawMiniAlignment` function.

#### Sequence Logos

CIAlign can generate traditional sequence logos [26] or sequence logos using rectangles instead of letters to show the information and base / amino acid content at each position, which can increase readability in less conserved regions.

### Interpretation

Some additional functions are provided to further interpret the alignment, for example plotting the number of sequences with non-gap residues at each position (the coverage), calculating a pairwise similarity matrix, and generating a consensus sequence with various options.

Given the toy example shown in Fig 1A, running all possible cleaning functions will lead to the markup plot shown in Fig 3A and the result shown in Fig 3B. In the markup plot each removed part is highlighted in a different colour corresponding to the function with which it was removed.

### Example Alignments

Four example alignments are provided within the software directory to demonstrate the functionality of CIAlign. Examples 1 and 2 use simulated sequences, examples 3 and 4 use real biological sequences and are designed to resemble the type of complex alignment many researchers encounter.

Example 1 is a very short alignment of six sequences which was generated manually by creating arbitrary sequences of nucleotides that would show every cleaning function while being as short as possible. This alignment contains an insertion, gaps at the ends of sequences, a very short sequence and some highly divergent sequences.

Example 2 is a larger alignment based on randomly generated amino acid sequences using RandSeq (a tool from ExPASy [27]) with an average amino acid composition, which were aligned with MAFFT v7.407, under the default settings [4]. The sequences were adjusted manually to reflect an alignment that would fully demonstrate the functionalities of CIAlign. It consists of many sequences that align well, however there are again a few problems: one sequence has a large insertion, one is very short, one is extremely divergent, and some have multiple gaps at the start and at the end. For Example 3, putative mitochondrial gene cytochrome C oxidase I (COI) sequences were identified by applying TBLASTN v2.9.0 [28] to the human COI sequence (GenBank accession NC_012920.1, positions 5,904–7,445, translated to amino acids), querying against 1,565 transcriptomic datasets from the NCBI transcriptome shotgun assembly (TSA) database [29] under the default settings. 2,855 putative COI transcripts were reverse complemented where required, and those corresponding to the COI gene of the primary host of the TSA dataset were identified using the BOLD online specimen identification engine [30] (accessed 07/10/2019) querying against the species level barcode records. The resulting 232 sequences were then aligned with MAFFT v7.407, under the default settings [4].

For Example 4, 91 sequences were selected from Example 3 to be representative of as many taxonomic families as possible and to exclude families with unclear phylogeny in the literature. These sequences were aligned with MAFFT v7.407 under the default settings and the alignment was refined with 1000 iterations. Robinson-Foulds distances of the resulting trees were calculated using ete3 compare [31].

Materials and methods for benchmarking and for large-scale examples with biological data are provided as Supplementary Materials and Methods.

## Results and Discussion

Here an example is presented and the visualisation functions are used to illustrate the functionality of CIAlign. Results will differ when using different parameters and thresholds. CIAlign was applied to the Example 2 alignment with the following options:

~~~
python3 CIAlign.py --infile INFILE --outfile_stem OUTFILE_STEM –all
~~~

Using these settings on the alignment in Fig 4A results in the markup shown in Fig 4B and the output shown in Fig 4C. The markup shows which function has removed each sequence or position. The benefits of CIAlign are clear in this simulation – the single poorly aligned sequence, the large insertion, very short sequences, and gap-only columns have been removed, and the unreliably aligned end segments of the sequences have been cropped. The resulting alignment is significantly shorter, which will speed up and simplify any further analysis. The clear graphical representation makes it easy to see what has been removed, so in the case of over-trimming the user can intervene and adjust functions and parameters.

**Fig 4.**
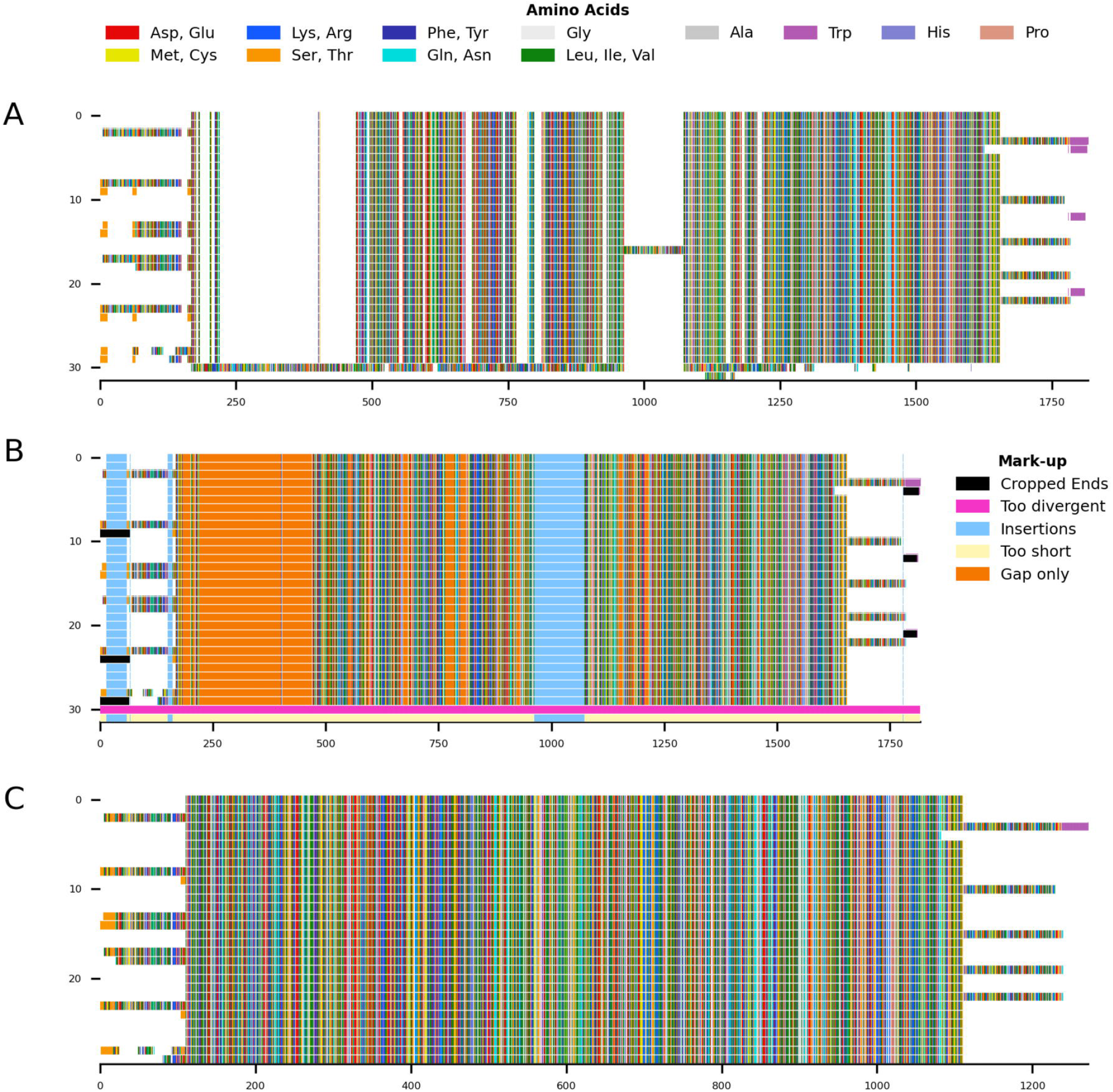
Mini alignments showing the main functionalities of CIAlign based on Example 2. **a** Input alignment before application of CIAlign, generated using the command “ `CIAlign --infile example2.fasta --plot_input`”. **b** Alignment markup showing areas that were removed by CIAlign, generated using the command “*CIAlign --infile example2*.*fasta --all*”. **c** Output alignment after application of CIAlign, generated using the command “`CIAlign --infile example2.fasta --all`”. Subplots were generated using the `drawMiniAlignment` function.

In order to demonstrate the use of CIAlign on real biological sequences, an alignment was generated based on the COI gene commonly used in phylogenetic analysis and DNA barcoding [30]. As CIAlign addresses some common problems encountered when generating an MSA based on *de novo* assembled transcripts, which tend to have a higher error rates at transcript ends, gaps due to difficult to assemble regions and divergent sequences due to chimeric connections between unrelated regions [11, 32], COI-like transcripts were identified by searching the NCBI transcriptome shotgun assembly database. Aligning these transcripts demonstrated several common problems – multiple insertions, poor alignment at the starts and ends of sequences, and a few divergent sequences resulting in excessive gaps (Fig 5A). This alignment was cleaned using the default CIAlign settings except the threshold for removing divergent sequences was reset to 50%, as some of the sequences are from evolutionarily distant species. Under these settings, CIAlign resolved several of the problems with the alignment: the insertions and highly divergent sequences were removed and the poorly aligned regions at the starts and ends of sequences were cropped (Fig 5B). One sequence and 6,029 positions were removed from the alignment and a total of 2,446 positions were cropped from the ends of 112 sequences. The processed alignment is 26.59% of the size of the input alignment. However, a minimal amount of actual sequence data (as opposed to gaps) was removed, with 85.70% of bases remaining.

**Fig 5.**
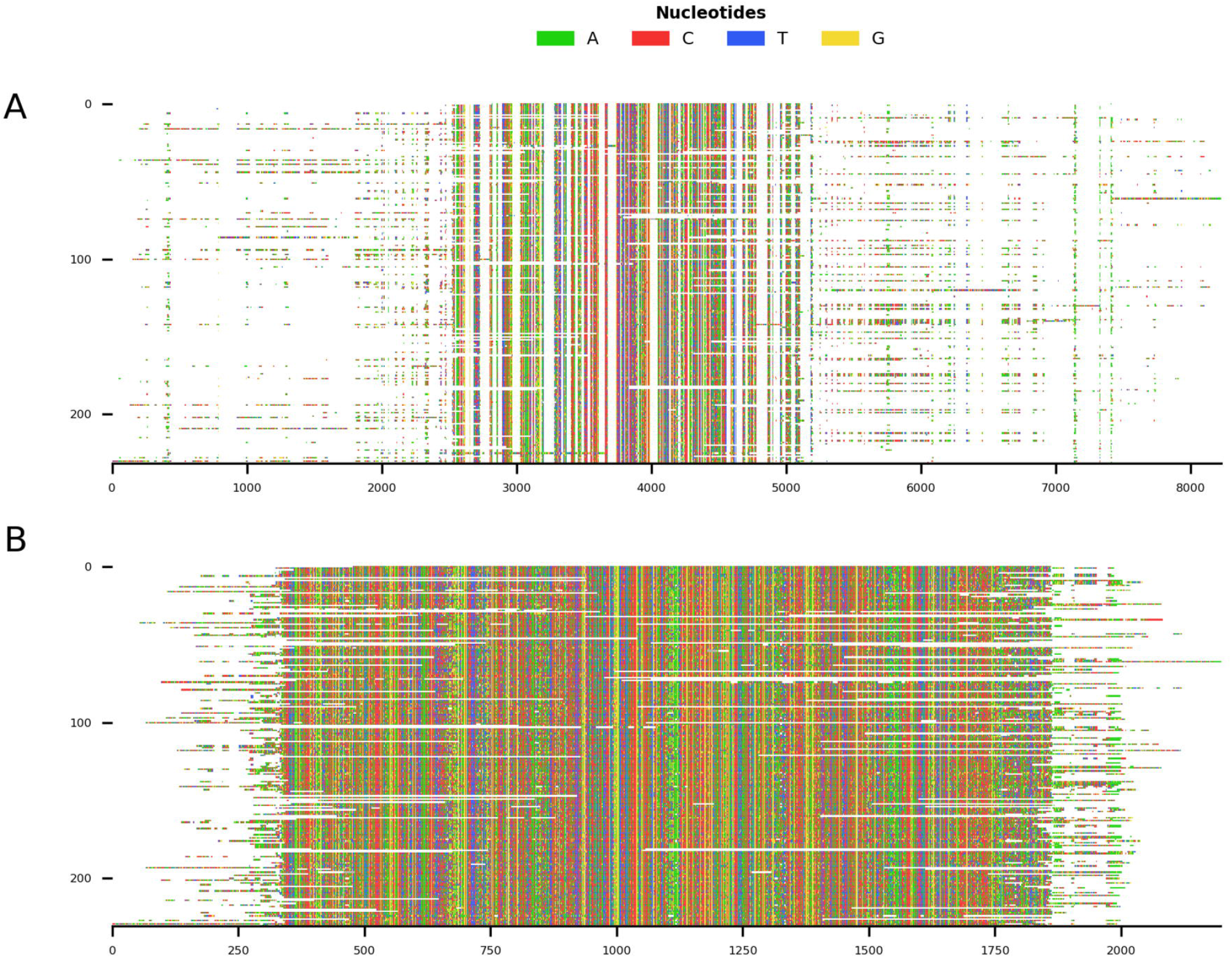
Mini alignments showing the main functionalities of CIAlign based on Example 3. **a** Input alignment before application of CIAlign, generated using the command “ `CIAlign --infile example3.fasta --plot_input”`. **b** Output alignment after application of CIAlign, generated using the command “`CIAlign --infile example3.fasta –all --remove_divergent_minperc 0.5`”. Subplots were generated using the`drawMiniAlignment` function.

A subset of this sequence set was selected to demonstrate the functionality of CIAlign in streamlining phylogenetic analysis. 91 COI-like transcripts from different taxonomic families of metazoa were selected from Example 3, incorporated into an MSA and cleaned using CIAlign with the same settings as above (Supp. Fig 1). 1,437 positions were removed from the alignment and a total of 289 positions were cropped from the ends of 17 sequences. The processed alignment is 70.67% of the size of the input alignment and 96.52% of bases remain. Phylogenetic trees were generated for the input alignment and for the alignment processed with CIAlign, using PhyML [33] under the GTR model plus the default settings. For the input alignment, PhyML used 138 MB of memory and took 532 seconds (on one Intel Core i7-7560U core with 4 GB of RAM, running at 2.40 GHz). For the cleaned alignment, on the same machine, PhyML used 109 MB of memory and took 243 seconds. The tree generated with the input alignment (Supp. Fig 1D) had a Robinson-Foulds [34] difference from a “correct” tree (generated manually based on the literature, Supp. Fig 1D) of 100.00 (normalised Robinson-Foulds 0.57, Quartet divergence [35] 0.159). The tree generated with the cleaned alignment (Supp. Fig 1E) had a Robinson-Foulds difference from the correct tree of 90.00 (normalised Robinson-Foulds 0.52, Quartet divergence 0.073) Therefore the tree based on the CIAlign cleaned alignment was generated more quickly, used less memory, and was more similar to the expected tree.

### Benchmarking with Simulated Data

We performed a series of benchmarking analyses on simulated data, in order to test and demonstrate the utility of the CIAlign cleaning functions, confirm the validity of our default parameter settings and ensure that running these functions does not have unexpected negative effects on downstream analyses.

First, CIAlign was tested using two tools (EvolvAGene [36], and INDELible [37]) which generate sets of unaligned sequences alongside “true” alignments and phylogenies expected to accurately represent the relationship between the sequences. We used these tools to determine if cleaning a user generated alignment with CIAlign affects its distance from the true alignment. Test alignments of the simulated data were created using four common alignment algorithms. These alignments were then cleaned with CIAlign with relaxed, moderate or stringent parameter settings (Supplementary Table 1). With relaxed CIAlign settings, a median of 0.19% of correct pairs of aligned residues (POARs) [38] were removed, for moderate settings 0.75% were removed and for stringent settings 3.76% (Fig 6A, Table 1). For comparison, the median total proportion of residues removed was 1.72% for relaxed, 2.40% for moderate and 3.86% for stringent (Fig 6A, Table 1). The median proportions of gap positions removed were much higher: 53-56% for all sets of parameters (Fig 6A, Table 1). This shows that with relaxed and moderate settings, running CIAlign has a very minimal impact on correctly aligned residues in the alignment, while a considerable amount of gaps and noise are removed. The more stringent settings should be used cautiously, however even with high stringency a large majority of correctly aligned residues remain and the majority of gaps are removed.

**Table 1.**
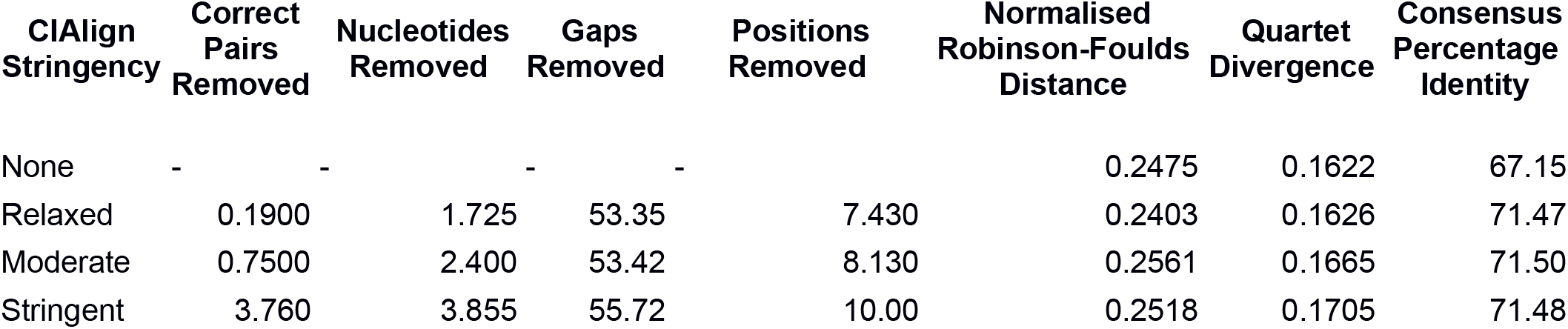
Table showing the effect of cleaning with CIAlign on simulated alignments.Correct pairs removed, nucleotides removed, gaps removed and positions removed are median percentage of the total in the test alignment which was removed by CIAlign. Normalised Robinson-Foulds distance and Quartet divergence are the mean proportion similarity between the benchmark tree and the tree generated based on the test alignment. Consensus percentage identity is the mean alignment similarity between the consensus and the EvolvAGene input sequence.

**Fig 6.**
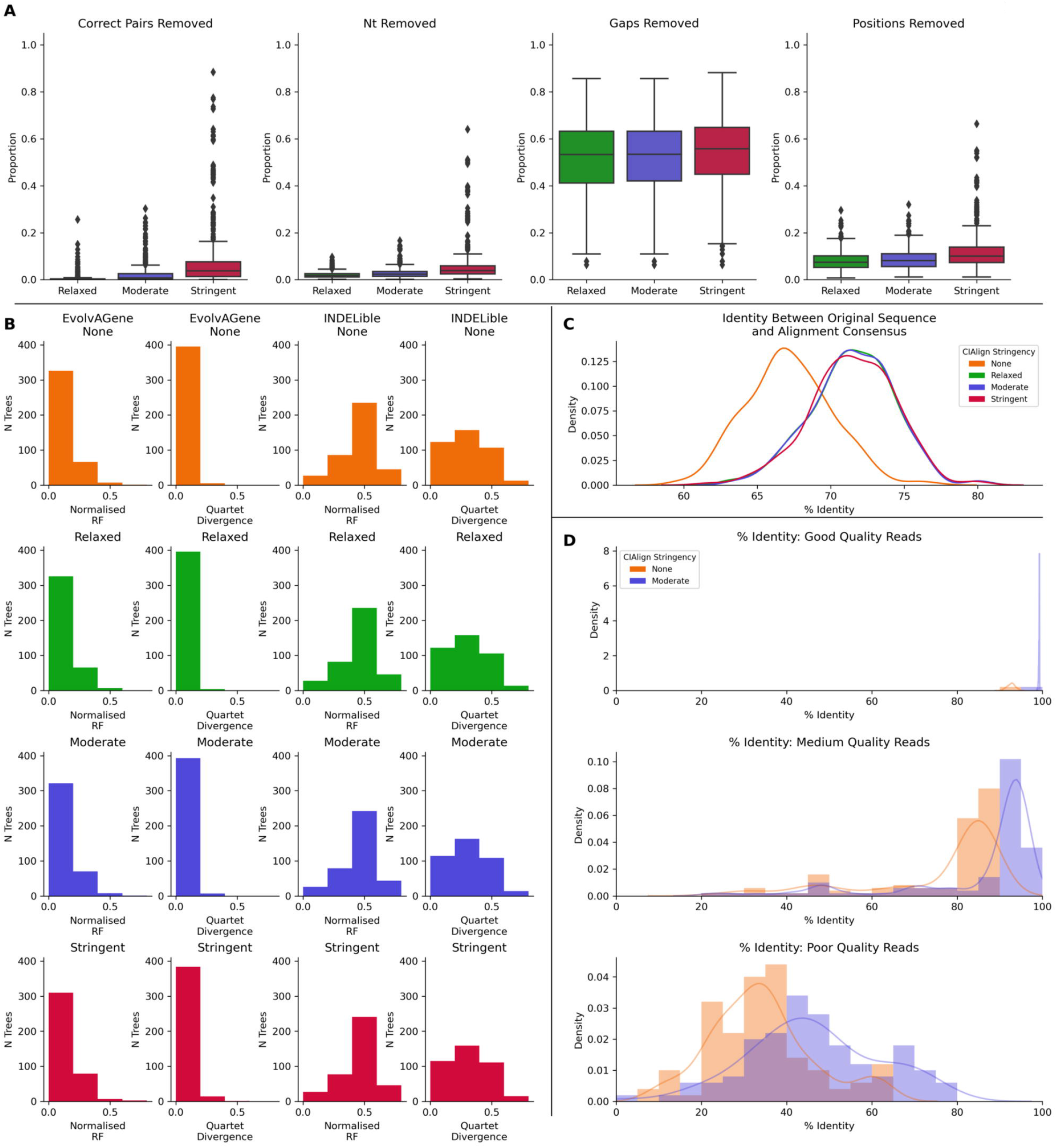
Metrics from benchmarking CIAlign with simulated data. **a** Box plots showing the impact of running CIAlign cleaning functions with relaxed (green, left box), moderate (blue, middle box) and stringent (red, right box) parameter values on alignments of sequences simulated using either EvolvAGene [36] or INDELible [37] (plots are combined for the two tools). From left to right, the y-axis represents proportion of correctly aligned pairs of residues [38] removed (identified by comparison with a benchmark alignment), proportion of total nucleotides (i.e. non-gap positions) removed, proportion of gaps removed, proportion of positions (gap or non-gap) removed. **b** Histograms showing the distribution of normalised Robinson-Foulds distances [34] and Quartet divergence [35] between benchmark trees and test trees without running CIAlign cleaning functions (orange) and after running CIAlign with the three sets of parameter values, for trees based on simulated sequences generated with EvolvAGene [36] (left two columns) and INDELible [37] (right two columns). **c** Density plot showing the distribution of the percentage identity between the input sequence to EvolvAGene [36] and a consensus sequence based on an alignment of the simulated sequences generated by this tool, without running CIAlign (orange) and after running CIAlign cleaning functions with the three sets of parameter values. **d** Density plots showing the distribution of the percentage identity between the input sequence to BadRead [39] and a consensus sequences generated with (blue) and without (orange) running CIAlign cleaning functions for alignments of good (top), medium (middle) and poor (bottom) quality simulated reads.

Phylogenetic trees were generated for each of these alignments to determine if cleaning with CIAlign impacts the distance between the true phylogenetic tree and a phylogenetic tree based on a test alignment (Fig 6B, Table 1. The mean normalised Robinson-Foulds distance and Quartet divergence [35] between the test trees and true trees were virtually unchanged by running CIAlign and none of the changes were statistically significant (p>0.05, Mann Whitney U test) (Fig 6B, Table 1).

We also compared the input sequence for our simulations to consensus sequences based on alignments with and without CIAlign cleaning. For all three stringency levels, CIAlign increased the percentage nucleotide identity between the consensus sequence and the input sequence by 4% to 5% (Fig 6C, Table 1). All of these changes are statistically significant (p<0.001, Mann-Whitney U test).

The long-read sequencing simulation tool BadRead [39] was used to demonstrate the use of CIAlign to remove common sources of error in long read sequencing data. Sequences were generated to represent low, moderate and high quality Oxford Nanopore reads based on an input genome, then aligned and cleaned with CIAlign with moderate settings (Supplementary Table 1). Using CIAlign increased the identity between the alignment consensus and the input sequence significantly for all read quality levels - by 6.57% for high quality reads, 9.51% for moderate quality reads and 12.25% for poor quality reads (Fig 6D) (all p<0.001, Mann-Whitney U test). For the high quality reads, the reads cleaned with CIAlign generated consensus sequences almost identical to the input sequence, with a mean of 99.24% identity (Fig 6D).

Full output tables for all three sets of simulations are available online at github.com/KatyBrown/CIAlign/benchmarking/tables and the simulated data and alignments at github.com/KatyBrown/benchmarking_data_CIAlign.

### Examples of Using CIAlign with Biological Data

We also used CIAlign to clean real biological data from several online databases, in order to test and demonstrate its usefulness in automated processing of different types of sequencing data.

#### Cleaning Pfam Alignments

The Pfam database provides manually curated seed alignments for over 17,000 protein families, plus much larger automatically generated full alignments containing sequences identified by database searching [40]. CIAlign cleaning functions were applied to seed and full alignments for 500 Pfam domains and consensus sequences were generated for both alignments, before and after cleaning. Randomly selected sequences from the full alignment were then compared to each consensus. For the full alignments, the mean identity between the consensus sequence and the alignment sequences increased by 10.71% (p < 0.001, Mann-Whitney U test) after cleaning with CIAlign (Fig 7A). For the seed alignments identity also increased significantly, by 4.89% (p < 0.001, Mann-Whitney U test) (Fig 7A). After running CIAlign, the full alignment consensus approaches the level of similarity to the alignment sequences which is seen for seed alignment consensus, despite the full alignment having undergone no manual curation (Fig 7A). Even for the curated seed alignments, cleaning with CIAlign further increases the similarity between the consensus and the aligned sequences.

**Fig 7.**
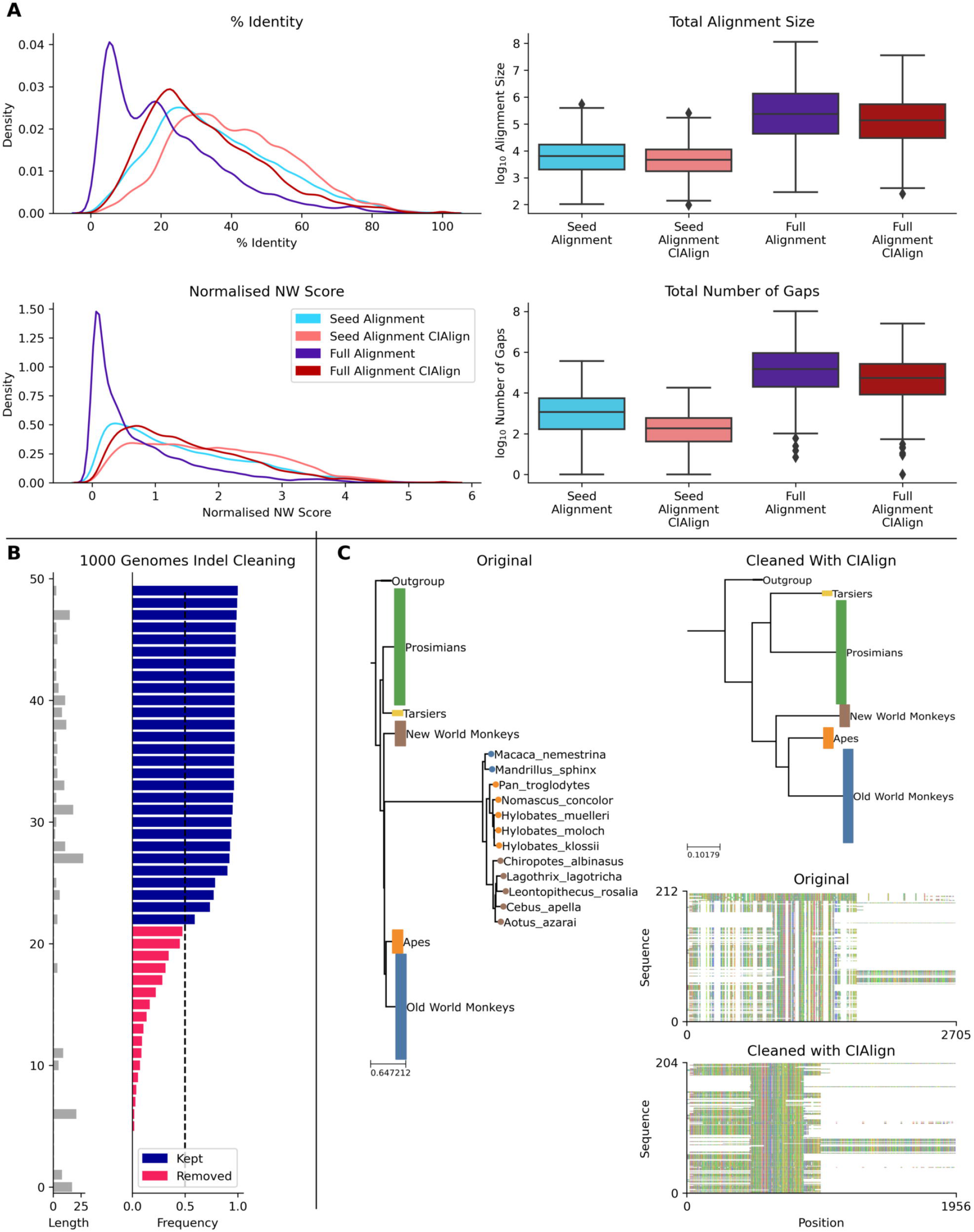
Metrics from using CIAlign with biological data. **a** Left, density plots showing the distribution of percentage identity (top) and normalised Needleman-Wunsch score (bottom) between samples of sequences from the Pfam [40] full alignments and consensus sequences generated based on Pfam seed alignments without (light blue) and with (light red) CIAlign cleaning and Pfam full alignments without (dark blue) and with (dark red) CIAlign cleaning. Right, box plots showing the alignment total size (top) and number of gaps (bottom) for these four alignments. **b** Left, bar chart showing the size of insertions from the 1000 genomes data [41] used to test the ability of CIAlign to remove insertions and deletions. Right, bar chart showing the proportion of sequences in which these insertions were present in data from 162 individuals and whether they were (pink) or were not (blue) removed by the CIAlign remove insertions function. **C** Left, phylogenetic tree based on an alignment of sequences from the 10k trees project [42] for the 12s ribosomal gene in primates. Colours represent known monophyletic groups of primates. Nodes have been collapsed where multiple sequences from the same group formed a monophyletic clade. Sequences annotated with circles were removed by CIAlign. Top-right, tree based on the same alignment after cleaning with CIAlign, which removed the outlying group. Bottom-right, mini alignments showing the effect of running CIAlign on this alignment.

#### Removing Insertions and Deletions from Human Genes

To demonstrate the ability of CIAlign to remove non-majority indels, we used data for 50 indels across over 150 individuals from the 1000 genomes project [41], which has annotated insertions and deletions for individual human genomes. In all cases, CIAlign removed all insertions present in a majority of samples and ignored all insertions present in a minority of samples (Fig 7B).

#### Removing Outliers

CIAlign can also be used to remove clear outliers from an alignment, for example prior to phylogenetic analysis. To illustrate this, we ran the CIAlign cleaning functions on data from the mammalian 10K trees project [42]. Three single-gene trees were identified with clear outliers, the 12S ribosomal gene from primates and the *APOB* and *RAG1* genes from Carnivora. The issues with these trees are shown in Fig 7C and Supp Fig 2. CIAlign successfully removed the outlying group, without removing any other sequences, in all three of these cases.

### Comparison with Other Software

While the functionality of CIAlign has some overlaps with other software, for example Jalview [43], Gblocks [7] and trimAl [8], the presented software can be seen as complementary to these, with some different features and applications. Jalview is designed for manual curation of alignments, but it is unsuitable for a simple overview of large alignments and does not provide the option of editing automatically, which is useful in large batch applications and ensures reproducibility. Gblocks is designed to choose blocks from an alignment that would be suitable for phylogenetic analysis, which is too restrictive for many other purposes. Some functionalities of trimAl overlap with those of CIAlign; however, trimAl is designed to algorithmically define and remove any poorly aligned regions whereas CIAlign is designed to remove specific MSA issues, as defined by the user, for different downstream applications. For highly divergent alignments, trimAl can be too sensitive and remove useful regions.

CIAlign also provides additional visualisation options. Therefore, CIAlign should be seen as a tool that aims to fill in the gaps that exist in currently available software.

### Parameters

Having as many parameters as possible to allow as much user control as possible gives greater flexibility. However, this also means that these parameters should be adjusted, which requires a good understanding of the cleaning functions and the MSA in question. CIAlign offers default parameters selected to be often applicable based on our benchmarking simulations and testing with different types of data. However, parameter choice highly depends on MSA divergence and the downstream application. To choose appropriate values it is recommended to first run CIAlign with all default parameters and then adjust these parameters based on the results. Since the mini alignments show what has been removed by which functions it is straightforward to identify the effect of each function and any changes to the parameters which may be required.

New features are in progress to be added in the future, such as collapsing very similar sequences, removing divergent columns, and making the colour scheme for the bases or amino acids customisable.

## Conclusion

CIAlign is a highly customisable tool which can be used to clean multiple sequence alignments and address several common alignment problems. Due to its multiple user options it can be used for many applications. CIAlign provides clear visual output showing which positions have been removed and for what reason, allowing the user to adjust the parameters accordingly. A number of additional visualisation and interpretation options are provided.

## Supporting information

Supplementary References

## Availability

**Current release, v1.0.10**: doi.org/10.5281/zenodo.4650727

(corresponds to github.com/KatyBrown/CIAlign/releases/tag/v1.0.10)

**GitHub**: github.com/KatyBrown/CIAlign

**pip3**: pypi.org/project/cialign

## Supplementary Information

**Supplementary Figure 1.**
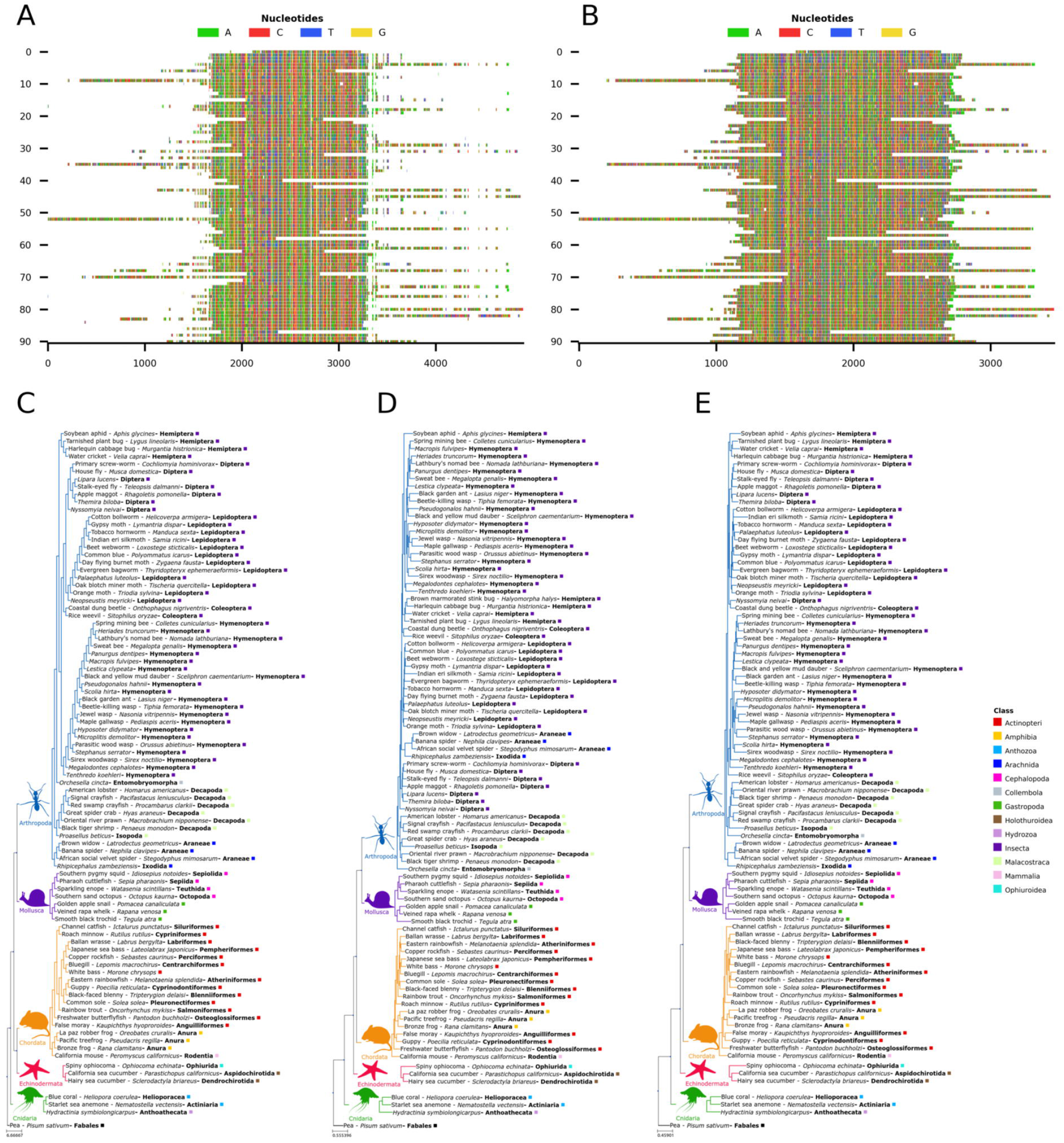
Mini alignments and phylogenetic trees showing the application of CIAlign to phylogenetic data, based on Example 4, a subset of Example 3. **a** Input alignment before application of CIAlign, generated using the command “*CIAlign --infile example4*.*fasta --plot_input”*. **b** Output alignment after application of CIAlign, generated using the command “*CIAlign --infile example4*.*fasta –all --remove_divergent_minperc 0*.*5*”. Subplots were generated using the “drawMiniAlignment function. **c** Phylogenetic tree generated manually using the literature to show the current best estimate for the phylogenetic relationships between these 91 families of metazoa. Relationships are based on the literature listed in the Supp. References. **d** PhyML phylogenetic tree generated under the GTR model plus default settings on the input alignment before application of CIAlign. **e** PhyML phylogenetic tree generated under the GTR model plus default settings on the cleaned alignment after application of CIAlign. In (c-e) branch colours correspond to the labelled phyla, coloured squares indicate class and bold text indicates order. Common names are shown where available.

**Supplementary Figure 2.**
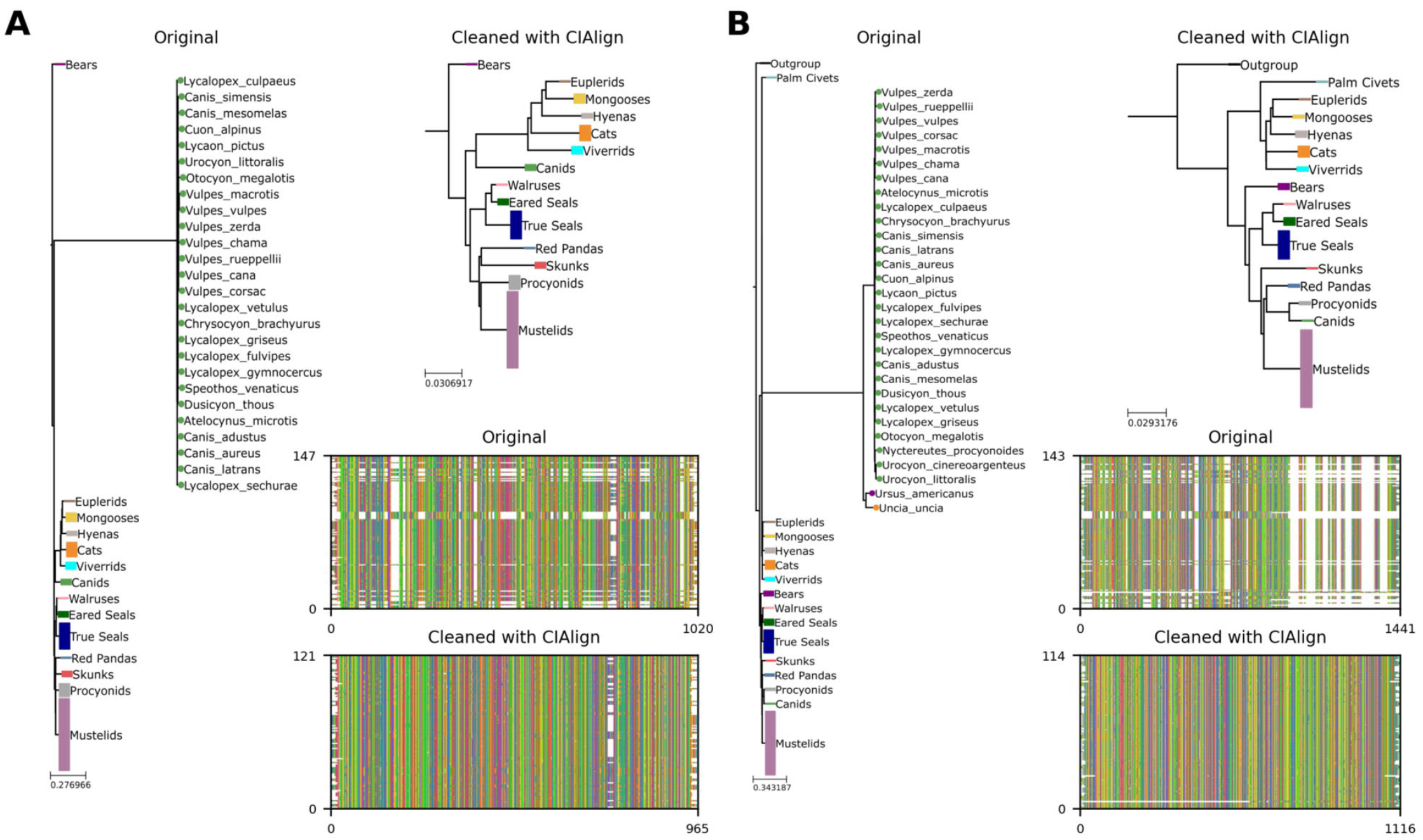
**a** Left, phylogenetic tree based on an alignment of sequences from the 10k trees project [42] for the *APOB* gene in Carnivora. Colours represent known monophyletic families of Carnivora. Nodes have been collapsed where multiple sequences from the same family formed a monophyletic clade. Sequences annotated with circles were removed by CIAlign. Top-right, tree based on the same alignment after cleaning with CIAlign, which removed the outlying group. Bottom-right, mini alignments showing the effect of running CIAlign on this alignment. **b** As for **a**, but for the *RAG1* gene in Carnivora.

**Supplementary Table 1.** Relaxed, moderate and stringent parameter settings used for benchmarking.

## References

1. Boswell, RD. Sequence alignment by word processor. Trends Biochem Sci. 1987;12:279–80.

2. Higgins DG, Sharp PM. CLUSTAL: a package for performing multiple sequence alignment on a microcomputer. Gene. 1988;73:237–44.

3. Edgar RC. MUSCLE: a multiple sequence alignment method with reduced time and space complexity. BMC Bioinformatics. 2004;5:113.

4. Katoh K, Misawa K, Kuma K, Miyata T. MAFFT: a novel method for rapid multiple sequence alignment based on fast Fourier transform. Nucleic Acids Res. 2002;30:3059–66.

5. Notredame C, Higgins DG, Heringa J. T-coffee: a novel method for fast and accurate multiple sequence alignment11Edited by J. Thornton. J Mol Biol. 2000;302:205–17.

6. Needleman SB, Wunsch CD. A general method applicable to the search for similarities in the amino acid sequence of two proteins. J Mol Biol. 1970;48:443–53.

7. Talavera G, Castresana J. Improvement of phylogenies after removing divergent and ambiguously aligned blocks from protein sequence alignments. Syst Biol. 2007;56:564–77.

8. Capella-Gutiérrez S, Silla-Martínez JM, Gabaldón T. trimAl: a tool for automated alignment trimming in large-scale phylogenetic analyses. Bioinformatics. 2009;25:1972–3.

9. Stamatakis A. RAxML version 8: a tool for phylogenetic analysis and post–analysis of large phylogenies. Bioinforma Oxf Engl. 2014;30:1312–3.

10. Kumar S, Stecher G, Li M, Knyaz C, Tamura K. MEGA X: Molecular Evolutionary Genetics Analysis across Computing Platforms. Mol Biol Evol. 2018;35:1547–9.

11. Bushmanova E, Antipov D, Lapidus A, Prjibelski AD. rnaSPAdes: a de novo transcriptome assembler and its application to RNA-Seq data. GigaScience. 2019;8. doi:10.1093/gigascience/giz100.

12. Richterich P. Estimation of Errors in “Raw” DNA Sequences: A Validation Study. Genome Res. 1998;8:251–9.

13. Tyler AD, Mataseje L, Urfano CJ, Schmidt L, Antonation KS, Mulvey MR, et al. Evaluation of Oxford Nanopore’s MinION Sequencing Device for Microbial Whole Genome Sequencing Applications. Sci Rep. 2018;8:1–12.

14. Magi A, Giusti B, Tattini L. Characterization of MinION nanopore data for resequencing analyses. Brief Bioinform. 2017;18:940–53.

15. Fitch WM, Smith TF. Optimal sequence alignments. PNAS. 1983;80:1382–6.

16. Tettelin H, Masignani V, Cieslewicz MJ, Donati C, Medini D, Ward NL, et al. Genome analysis of multiple pathogenic isolates of Streptococcus agalactiae: implications for the microbial “pan-genome.” Proc Natl Acad Sci U S A. 2005;102:13950–5.

17. Hu B, Xie G, Lo C-C, Starkenburg SR, Chain PSG. Pathogen comparative genomics in the next-generation sequencing era: genome alignments, pangenomics and metagenomics. Brief Funct Genomics. 2011;10:322–33.

18. Sayyari E, Whitfield JB, Mirarab S. Fragmentary Gene Sequences Negatively Impact Gene Tree and Species Tree Reconstruction. Mol Biol Evol. 2017;34:3279–91.

19. Schulz F, Roux S, Paez-Espino D, Jungbluth S, Walsh DA, Denef VJ, et al. Giant virus diversity and host interactions through global metagenomics. Nature. 2020;578:432–6.

20. Käfer S, Paraskevopoulou S, Zirkel F, Wieseke N, Donath A, Petersen M, et al. Re-assessing the diversity of negative strand RNA viruses in insects. PLoS Pathog. 2019;15. doi:10.1371/journal.ppat.1008224.

21. Bäckström D, Yutin N, Jørgensen SL, Dharamshi J, Homa F, Zaremba-Niedwiedzka K, et al. Virus Genomes from Deep Sea Sediments Expand the Ocean Megavirome and Support Independent Origins of Viral Gigantism. mBio. 2019;10. doi:10.1128/mBio.02497-18.

22. Petyuk VA, Gatto L, Payne SH. Reproducibility and Transparency by Design. Mol Cell Proteomics. 2019. doi:10.1074/mcp.IP119.001567.

23. Brito JJ, Li J, Moore JH, Greene CS, Nogoy NA, Garmire LX, et al. Recommendations to enhance rigor and reproducibility in biomedical research. 2020. https://arxiv.org/abs/2001.05127v2.

24. Langille MGI, Ravel J, Fricke WF. “Available upon request”: not good enough for microbiome data! Microbiome. 2018;6:8.

25. Hunter JD. Matplotlib: A 2D Graphics Environment. Comput Sci Eng. 2007;9:90–5.

26. Schneider TD, Stephens RM. Sequence logos: a new way to display consensus sequences. Nucleic Acids Res. 1990;18:6097–100.

27. Gasteiger E, Gattiker A, Hoogland C, Ivanyi I, Appel RD, Bairoch A. ExPASy: The proteomics server for in-depth protein knowledge and analysis. Nucleic Acids Res. 2003;31:3784–8.

28. Camacho C, Coulouris G, Avagyan V, Ma N, Papadopoulos J, Bealer K, et al. BLAST+: architecture and applications. BMC Bioinformatics. 2009;10:421.

29. Transcriptome Shotgun Assembly Sequence Database. National Center for Biotechnology Information, Bethesda, Maryland, USA. 2012. https://www.ncbi.nlm.nih.gov/genbank/tsa/. Accessed 08 Oct 2019.

30. bold: The Barcode of Life Data System (http://www.barcodinglife.org). https://www.ncbi.nlm.nih.gov/pmc/articles/PMC1890991/. Accessed 6 Apr 2020.

31. Huerta-Cepas J, Serra F, Bork P. ETE 3: Reconstruction, Analysis, and Visualization of Phylogenomic Data. Mol Biol Evol. 2016;33:1635–8.

32. Liao X, Li M, Zou Y, Wu F-X, Yi-Pan, Wang J. Current challenges and solutions of de novo assembly. Quant Biol. 2019;7:90–109.

33. Guindon S, Gascuel O. A simple, fast, and accurate algorithm to estimate large phylogenies by maximum likelihood. Syst Biol. 2003;52:696–704.

34. Robinson DF, Foulds LR. Comparison of phylogenetic trees. Math Biosci. 1981;53:131–47.

35. Smith MR. Bayesian and parsimony approaches reconstruct informative trees from simulated morphological datasets. Biol Lett. 2019;15:20180632.

36. Hall BG. Simulating DNA coding sequence evolution with EvolveAGene 3. Mol Biol Evol. 2008;25:688–95.

37. Fletcher W, Yang Z. INDELible: a flexible simulator of biological sequence evolution. Mol Biol Evol. 2009;26:1879–88.

38. Thompson JD, Plewniak F, Poch O. A comprehensive comparison of multiple sequence alignment programs. Nucleic Acids Res. 1999;27:2682–90.

39. Wick RR. Badread: simulation of error-prone long reads. J Open Source Softw. 2019;4:1316.

40. Finn RD, Bateman A, Clements J, Coggill P, Eberhardt RY, Eddy SR, et al. Pfam: the protein families database. Nucleic Acids Res. 2014;42 Database issue:D222–30.

41. Auton A, Abecasis GR, Altshuler DM, Durbin RM, Abecasis GR, Bentley DR, et al. A global reference for human genetic variation. Nature. 2015;526:68–74.

42. Arnold C, Matthews LJ, Nunn CL. The 10kTrees website: A new online resource for primate phylogeny. Evol Anthropol Issues News Rev. 2010;19:114–8.

43. Waterhouse AM, Procter JB, Martin DMA, Clamp M, Barton GJ. Jalview Version 2--a multiple sequence alignment editor and analysis workbench. Bioinforma Oxf Engl. 2009;25:1189–91.

