## Supplementary Materials and Methods for "CIAlign - A highly customisable command line tool to clean, interpret and visualise multiple sequence alignments"

### Benchmarking

*Sequence Simulation*

Simulated sequences and alignments were generated using EvolveAGene4 [1], INDELible v1.0.3 [2] and BadRead v0.1.5 [3]. For EvolveAGene, 100 simulations were performed based on the human GAPDH coding sequence (Genbank accession: NC_000012.12, CCDS8549.1) with eight taxa, using the default settings and an average branch length of 0.62 (as recommended in the manual). For each simulation, this tool generated eight simulated coding sequences with associated true nucleotide and amino acid alignments and phylogenetic trees. For INDELible 100 simulations were performed using the 'GTRexample' parameters from the manual and the 'nucleotide 1' algorithm to simulate nucleotide evolution under the GTR model. The guide tree used and control files are available on the CIAlign GitHub. BadRead sequences were generated for 100 simulations each for the settings proposed for "very nice", "mediocre" and "very bad" Oxford Nanopore reads in the BadRead documentation. All BadRead simulations were performed on the full length deformed wing virus reference sequence, Genbank Accession NC_004830.2. Shell scripts used to run these simulations are available in the benchmarking directory for each tool on the CIAlign GitHub as run_simulations.sh. The output data from all simulations is available at github.com/KatyBrown/benchmarking_data_CIAlign.

*Alignment*

The EvolveAGene and INDELible sequences were aligned using Clustal (Omega version 1.2.4, [4]), with the default parameters plus --auto, MUSCLE (version 3.8.31, [5]), with the default parameters plus 100 iterations and MAFFT (version 7.464, [6]) with the default parameters (global) and with default parameters plus –localpair (local), both with 1000 iterations. BadRead sequences are expected to be incomplete, so were aligned only with MAFFT with the default parameters plus –localpair. Shell scripts used to generate these alignments are available in the benchmarking directory for each tool on the CIAlign GitHub as run_alignment.sh). All alignments are available at github.com/KatyBrown/benchmarking_data_CIAlign.

*CIAlign Cleaning*

CIAlign was used to clean all alignments with relaxed, moderate and high stringency parameters, parameter values are listed in Supplementary Table 1. Generally the default parameters were used as the moderate parameter settings, the exception to this is remove_divergent, which was used at lower stringency for INDELible and BadRead because these sequences are very divergent. Shell scripts used to run CIAlign are available in the benchmarking directory for each tool on the CIAlign GitHub as run_cialign.sh. All cleaned alignments are available at github.com/KatyBrown/benchmarking_data_CIAlign.

*Comparisons*

Correctly aligned sequence pairs were identified by comparison with the benchmark alignments generated by the software and calculated using the get_POARs function implemented in the AlignmentStats module in the CIAlign benchmarking functions directory. Consensus sequences in all cases were generated in CIAlign with consensus type majority_nongap and phylogenetic trees with FastTree2 with a GTR model (v2.1.10, [7]). Identity and Needleman-Wunsch scores between pairs of sequences were calculated using the Needle tool from the EMBOSS package (v6.5.7.0, [8]). Robinson-Foulds distances were calculated using the compare function of the Python package ete3 (v3.1.1, [9]). Quartet divergence was calculated using the tqdist algorithm [10] implemented in the R package Quartet [11] [12], with similarity=FALSE to calculate divergence rather than similarity. All consensus sequences and trees are available at github.com/KatyBrown/benchmarking_data_CIAlign. All scores are available in Supplementary Table 1 (for EvolvAGene and INDELible) and Supplementary Table 2 (for BadRead) on the CIAlign GitHub in the benchmarking/tables directory.

**Biological Data**

*Cleaning Pfam Alignments*

A random sample of 500 Pfam domains was selected (listed in full in the online Supplementary Table 3 in the benchmarking/tables directory) and seed and full alignments for each domain were downloaded from Pfam release 34.0 [13]. All alignments were cleaned with CIAlign remove_insertions (insertion_min_size 1 and insertion_max_size 500) and crop_ends (crop_ends_mingap_perc 0.01, crop_ends_redefine_perc 0.1). Consensus sequences were generated in CIAlign with consensus type majority_nongap. 250 sequences were selected at random from each full alignment and aligned to consensus sequences from the seed and full alignments before and after CIAlign cleaning using the Needle tool from the EMBOSS package (v6.5.7.0, [8]), this tool was also using to calculate identity scores and Needleman-Wunsch scores.

*Removing Insertions and Deletions from Human Genes*

Human protein coding gene positions were identified based on human genome build GRCh38.p13 from Ensembl release 103 [14]. Genes were selected at random and single gene VCF files downloaded from the human genome project release 20170504 [15] via Tabix (v 1.1.9, [16]). Genes with at least one indel variant were selected until 25 insertions and 25 deletions (relative to the reference) had been identified, covering 32 genes in total. Individual genomes were then selected at random until a non-reference call for each variant had been identified in at least one individual. This process gave a set of 162 individuals. Separate VCF files were generated for each individual for each of the 32 genes and converted to FASTA files using bcftools (v1.9 [17]) with the -H A option to prioritise alternate variants. These FASTA files were aligned with MAFFT (version 7.464, [6]) with the default settings. They were then cleaned with CIAlign remove_insertions with insertion_min_size 1 and insertion_max_size 50. Full results including all gene Ensembl IDs, variant IDs and sample IDs are available in the online Supplementary Table 4 on the CIAlign GitHub in the benchmarking/tables directory.

*Removing Outliers*

Unaligned single gene FASTA files were downloaded for mammalian genes from the 10k trees project [18] (version 3 for primates, version 1 for Carnivora). These files were aligned MAFFT (version 7.464, [6]) with the default settings. The alignments were then cleaned with the CIAlign remove insertions, crop ends, remove short and remove divergent functions, with default settings except for remove_divergent_minperc, which was set to 0.7 (as these are quite conserved sequences). Alignments before and after cleaning were visualised as CIAlign mini alignments with the default settings. Phylogenetic trees were generated for alignments before and after cleaning using FastTree2 with a GTR model (v2.1.10, [7]). Trees were pruned and labelled using the ete3 Python package (v3.1.1 [9]).

**References**

1. Hall BG. Simulating DNA coding sequence evolution with EvolveAGene 3. Mol Biol Evol. 2008;25:688–95.

2. Fletcher W, Yang Z. INDELible: a flexible simulator of biological sequence evolution. Mol Biol Evol. 2009;26:1879–88.
3. Wick RR. Badread: simulation of error-prone long reads. J Open Source Softw. 2019;4:1316.
4. Higgins DG, Sharp PM. CLUSTAL: a package for performing multiple sequence alignment on a microcomputer. Gene. 1988;73:237–44.
5. Edgar RC. MUSCLE: a multiple sequence alignment method with reduced time and space complexity. BMC Bioinformatics. 2004;5:113.
6. Katoh K, Standley DM. MAFFT multiple sequence alignment software version 7: improvements in performance and usability. Mol Biol Evol. 2013;30:772–80.
7. Price MN, Dehal PS, Arkin AP. FastTree 2--approximately maximum-likelihood trees for large alignments. PloS One. 2010;5:e9490.
8. Rice P, Longden I, Bleasby A. EMBOSS: the European Molecular Biology Open Software Suite. Trends Genet TIG. 2000;16:276–7.
9. Huerta-Cepas J, Serra F, Bork P. ETE 3: Reconstruction, Analysis, and Visualization of Phylogenomic Data. Mol Biol Evol. 2016;33:1635–8.
10. Sand A, Holt MK, Johansen J, Brodal GS, Mailund T, Pedersen CNS. tqDist: a library for computing the quartet and triplet distances between binary or general trees. Bioinformatics. 2014;30:2079–80.
11. Smith MR. Quartet: comparison of phylogenetic trees using quartet and split measures. 2019.
12. Smith MR. Bayesian and parsimony approaches reconstruct informative trees from simulated morphological datasets. Biol Lett. 2019;15:20180632.
13. Finn RD, Bateman A, Clements J, Coggill P, Eberhardt RY, Eddy SR, et al. Pfam: the protein families database. Nucleic Acids Res. 2014;42 Database issue:D222–30.
14. Yates AD, Achuthan P, Akanni W, Allen J, Allen J, Alvarez-Jarreta J, et al. Ensembl 2020. Nucleic Acids Res. 2020;48:D682–8.
15. Auton A, Abecasis GR, Altshuler DM, Durbin RM, Abecasis GR, Bentley DR, et al. A global reference for human genetic variation. Nature. 2015;526:68–74.
16. Li H. Tabix: fast retrieval of sequence features from generic TAB-delimited files. Bioinformatics. 2011;27:718–9.
17. Danecek P, Bonfield JK, Liddle J, Marshall J, Ohan V, Pollard MO, et al. Twelve years of SAMtools and BCFtools. GigaScience. 2021;10. doi:10.1093/gigascience/giab008.

18. Arnold C, Matthews LJ, Nunn CL. The 10kTrees website: A new online resource for primate phylogeny. Evol Anthropol Issues News Rev. 2010;19:114–8.
