## Supplementary References for "CIAlign - A highly customisable command line tool to clean, interpret and visualise multiple sequence alignments"

1. Fong JJ, Brown JM, Fujita MK, Boussau B. A Phylogenomic Approach to Vertebrate Phylogeny Supports a Turtle-Archosaur Affinity and a Possible Paraphyletic Lissamphibia. PLoS One. 2012;7. doi:10.1371/journal.pone.0048990.

2. Wolfe JM, Breinholt JW, Crandall KA, Lemmon AR, Lemmon EM, Timm LE, et al. A phylogenomic framework, evolutionary timeline and genomic resources for comparative studies of decapod crustaceans. Proceedings of the Royal Society B: Biological Sciences. 2019;286:20190079.

3. Almeida D, Maldonado E, Vasconcelos V, Antunes A. Adaptation of the Mitochondrial Genome in Cephalopods: Enhancing Proton Translocation Channels and the Subunit Interactions. PLOS ONE. 2015;10:e0135405.

4. Regier JC, Shultz JW, Zwick A, Hussey A, Ball B, Wetzer R, et al. Arthropod relationships revealed by phylogenomic analysis of nuclear protein-coding sequences. Nature. 2010;463:1079–83.

5. Hughes LC, Ortí G, Huang Y, Sun Y, Baldwin CC, Thompson AW, et al. Comprehensive phylogeny of ray-finned fishes (Actinopterygii) based on transcriptomic and genomic data. Proc Natl Acad Sci USA. 2018;115:6249–54.

6. Wiegmann BM, Trautwein MD, Winkler IS, Barr NB, Kim J-W, Lambkin C, et al. Episodic radiations in the fly tree of life. PNAS. 2011;108:5690–5.

7. Li H, Leavengood JM, Chapman EG, Burkhardt D, Song F, Jiang P, et al. Mitochondrial phylogenomics of Hemiptera reveals adaptive innovations driving the diversification of true bugs. Proc Biol Sci. 2017;284. doi:10.1098/rspb.2017.1223.

8. Misof B, Liu S, Meusemann K, Peters RS, Donath A, Mayer C, et al. Phylogenomics resolves the timing and pattern of insect evolution. Science. 2014;346:763–7.

9. Feng Y-J, Blackburn DC, Liang D, Hillis DM, Wake DB, Cannatella DC, et al. Phylogenomics reveals rapid, simultaneous diversification of three major clades of Gondwanan frogs at the Cretaceous–Paleogene boundary. PNAS. 2017;114:E5864–70.

10. Kawahara AY, Plotkin D, Espeland M, Meusemann K, Toussaint EFA, Donath A, et al. Phylogenomics reveals the evolutionary timing and pattern of butterflies and moths. PNAS. 2019;116:22657–63.

11. Laumer CE, Fernández R, Lemer S, Combosch D, Kocot KM, Riesgo A, et al. Revisiting metazoan phylogeny with genomic sampling of all phyla. Proceedings of the Royal Society B: Biological Sciences. 2019;286:20190831.

12. Garrison NL, Rodriguez J, Agnarsson I, Coddington JA, Griswold CE, Hamilton CA, et al. Spider phylogenomics: untangling the Spider Tree of Life. PeerJ. 2016;4:e1719.

13. Gu X-B, Liu G-H, Song H-Q, Liu T-Y, Yang G-Y, Zhu X-Q. The complete mitochondrial genome of the scab mite Psoroptes cuniculi (Arthropoda: Arachnida) provides insights into Acari phylogeny. Parasites & Vectors. 2014;7:340.

14. McKenna DD, Shin S, Ahrens D, Balke M, Beza-Beza C, Clarke DJ, et al. The evolution and genomic basis of beetle diversity. PNAS. 2019;116:24729–37.

15. Peters RS, Meyer B, Krogmann L, Borner J, Meusemann K, Schütte K, et al. The taming of an impossible child: a standardized all-in approach to the phylogeny of Hymenoptera using public database sequences. BMC Biology. 2011;9:55.
